# Multivariable G-E interplay in the prediction of educational achievement

**DOI:** 10.1101/865360

**Authors:** A.G. Allegrini, V. Karhunen, J. R. I. Coleman, S. Selzam, K. Rimfeld, S. von Stumm, J.-B. Pingault, R. Plomin

## Abstract

Polygenic scores are increasingly powerful predictors of educational achievement. It is unclear, however, how sets of polygenic scores, which partly capture environmental effects, perform jointly with sets of environmental measures, which are themselves heritable, in prediction models of educational achievement.

Here, for the first time, we systematically investigate gene-environment correlation (rGE) and interaction (GxE) in the joint analysis of multiple genome-wide polygenic scores (GPS) and multiple environmental measures as they predict tested educational achievement (EA). We predict EA in a representative sample of 7,026 16-year-olds, with 20 GPS for psychiatric, cognitive and anthropometric traits, and 13 environments (including life events, home environment, and SES) measured earlier in life. Environmental and GPS predictors were modelled, separately and jointly, in penalized regression models with out-of-sample comparisons of prediction accuracy, considering the implications that their interplay had on model performance.

Jointly modelling multiple GPS and environmental factors significantly improved prediction of EA, with cognitive-related GPS adding unique independent information beyond SES, home environment and life events. We found evidence for rGE underlying variation in EA (rGE = .36; 95% CIs = .29, .43). We estimated that 38% (95% CIs = 29%, 49%) of the GPS effects on EA were mediated by environmental effects, and in turn that 18% (95% CIs =12%, 25%) of environmental effects were accounted for by the GPS model. Lastly, we did not find evidence that GxE effects collectively contributed to multivariable prediction.

Our multivariable polygenic and environmental prediction model suggests widespread rGE and unsystematic GxE contributions to EA in adolescence.

## Introduction

Education is compulsory in nearly all countries because it provides children with the skills, such as literacy and numeracy, that are essential for successfully participating in society. How well children perform at school, indicated by their educational achievement (EA), predicts many important life outcomes, especially further education and occupational status (1). Quantitative genetic research based on twin studies showed that EA is 60% heritable throughout the school years (2, 3). These studies also suggested that about 20% of the variance of EA and other learning-related traits can be ascribed to shared environmental factors, for example growing up in the same family and going to the same school. However, the picture became more complicated with the discovery that ostensible measures of the environment associated with educational achievement showed genetic influence – most notably, parents’ educational attainment, socio-economic status (SES) and aspects of the home environment (4).

Quantitative genetic theory distinguishes two types of interplay between genetic and environmental effects, genotype-environment correlation (rGE) and genotype-environment interaction (GxE) (5). rGE occurs when an individual’s genotype covaries with environmental exposures. There are three types of rGE: passive, active and evocative. Passive rGE results from the inheritance of both genetic propensities and environments linked to parental genotypes. That is, individuals inherit from parents a genetic predisposition to a particular trait, but parental genotypes are also associated with rearing environments that, in turn, increase the likelihood of developing a particular trait. For example, individuals with stronger genetic predispositions to educational attainment tend to grow up in higher socioeconomic status families (6). Evocative rGE happens when individuals’ genetic propensities evoke a response from the surrounding environment; for example children’s predisposition to higher food intake might elicit restrictive food behaviors from their parents (7). Active rGE results from individuals actively selecting environments that are linked to their genetic propensity; for example, individuals with a higher genetic predisposition to educational attainment tend to migrate to economically prosperous regions that offer greater educational opportunities (8).

GxE, on the other hand, refers to genetic moderation of environmental effects. That is, when the effects of environmental exposures on phenotypes depend on individuals’ genotypes. Equivalently, environmentally moderated genetic effects occur when genetic effects on a phenotype depend on environmental exposures. Importantly, however, rGE may confound GxE effects (9). For example, if a genetic predisposition for a particular trait is found in a particular environment, it is difficult to know whether this represents rGE between the trait and the environment or true GxE. As before, this picture becomes even more complicated when we consider that environments are themselves heritable.

Research on GxE was rejuvenated when it became possible to include measured genetic and environmental factors in statistical models. Hundreds of studies were published purporting to show interactions between candidate genes and environmental measures as they predict behavioural traits. For example, a seminal GxE study in the field (10) showed that carriers of two copies of the short serotonine allele on the 5HTT gene exposed to adversity had an increased the risk for depression compared to their genetic counterpart. However, GxE effects such as these have a poor replication history (11, 12). The main problem with this approach is that it ignores the high polygenicity of complex traits, with a reductionist focus on single ‘candidate’ variants. This combined with typically small sample sizes, underpowered to detect the very small effects that can be expected for GxE, lead to a failure to replicate (13).

In complex traits, very few individual variants capture more than a tiny fraction of trait variance (14). Genome-wide polygenic scores (GPS) are the missing piece for investigating the interplay between genes and environment because they can theoretically capture genetic influences up to the limit of SNP-based heritability, which is usually 25-50% of the total heritability for behavioural traits. GPS are indices of an individual’s genetic propensity for a trait and are typically derived as the sum of the total number of trait-associated alleles across the genome, weighted by their respective association effect size estimated through genomewide association analysis (15). A GPS derived from a genome-wide association study of educational attainment (years of schooling) (16) predicts up to 15% of the variance of EA (17). As more powerful GPS become available, they have begun to be used widely in research on GxE (18–23) and rGE (7, 24–27).

Recently it has been possible to dissect the role of parental genetics on child achievement by splitting the parental genome into transmitted alleles (indexing passive rGE) and nontransmitted alleles (indexing environmentally transmitted parental genetic effects). The latter demonstrated that parental genotypes are associated with the environment they provide for the child (28, 29). In fact, a growing body of evidence is showing the importance of considering gene-environment correlation when assessing polygenic effects on trait variation (30, 31), especially for educationally relevant traits. Paralleling quantitative genetics results, a key point is that environmental measures are themselves heritable and GPS effects can be mediated by the environment, while environmental effects can be accounted for by genetics (genetic confounding). In this sense, polygenic scores for cognitive traits are not pure measures of genetic predisposition: their predictive power also captures environmental effects. For the same reason, environmental measures are not pure measures of the environment.

Rather than examining rGE and GxE for single polygenic scores and environmental measures, here we look at sets of GPS (32) and environmental measures. A multivariable approach is especially warranted for EA because twin analyses show that the high heritability (60%) of EA reflects many genetically influenced traits, including personality and behaviour problems in addition to cognitive traits (33, 34). Correspondingly, EA GPS is associated with a wide range of traits, including psychiatric, anthropometric and behavioural traits (35). Similarly, environmental predictors of EA are also intercorrelated (e.g. SES and home environment). However, it is not yet clear how sets of polygenic scores, partly capturing environmental effects, perform jointly with sets of environmental measures, which are themselves heritable, and the effect that their interplay (rGE and GxE) might have on prediction.

Here for the first time we systematically investigate the interplay of GPS and environmental measures in the multivariable prediction of tested educational achievement. We jointly analyse multiple GPS and multiple environmental measures, considering the effect of their interplay in out-of-sample prediction. Specifically, we test the joint prediction of 20 well-powered GPS for psychiatric, cognitive and anthropometric traits and 13 proximal and distal measured environments including life events, home environment and SES (see methods for descriptions of all measures). First, we model polygenic scores (henceforth G model) and environmental measures (henceforth E model), separately and jointly (full model), to predict educational achievement in penalized regression models (36) with out-of-sample tests of prediction accuracy. To investigate the relative contributions of the employed predictors to the full model, we carry out post-selection estimation (37) of partial regression coefficients, testing independent effects of single GPS and environmental measures. Second, we separate direct from mediated effects of the multivariable G and E models on EA and assess rGE defined in terms of the GPS and environmental measures employed. Finally, we assess GxE using a hierarchical group-lasso technique (38) to systematically discover two-way interactions between all GPS and environmental measures, and test their improvement in prediction of EA.

## Results

### Joint modelling of GPS and environmental effects

We tested three models for association with EA: all genetic factors (polygenic scores; G model), all environmental factors (measured environments; E model), and a joint model of all factors (full model). Joint modelling of both GPS and environmental measures achieved the best out-of-sample prediction compared to the G or E models considered separately. The full model predicted 37% of the variance (95% CI = 31.2, 43.1) in EA (Figure 1 panel A, Supplementary Table S2), an improvement of 7% in prediction compared to the E model (30.6%; 95% CI = 24.5, 36.6; Supplementary figure S1) and up to 19% improvement compared to the G model (18.8%; 95% CI = 13.5, 25.1; Supplementary Figure S2). Nested comparisons of the full model vs the G and E models separately suggested that the difference in out-of-sample prediction accuracy between models (Figure 1 panel B, Supplementary Table S2b) was significant for both the full model vs E (R^2^ median diff = 6.4%; 95% CI = 3.4, 9.7) and the full model vs G (R^2^ median diff = 18.16%; CI = 13.0, 23.3). Next, we untangled the specific independent contributions of GPS and measured environments to variation in EA.

**Figure 1.**
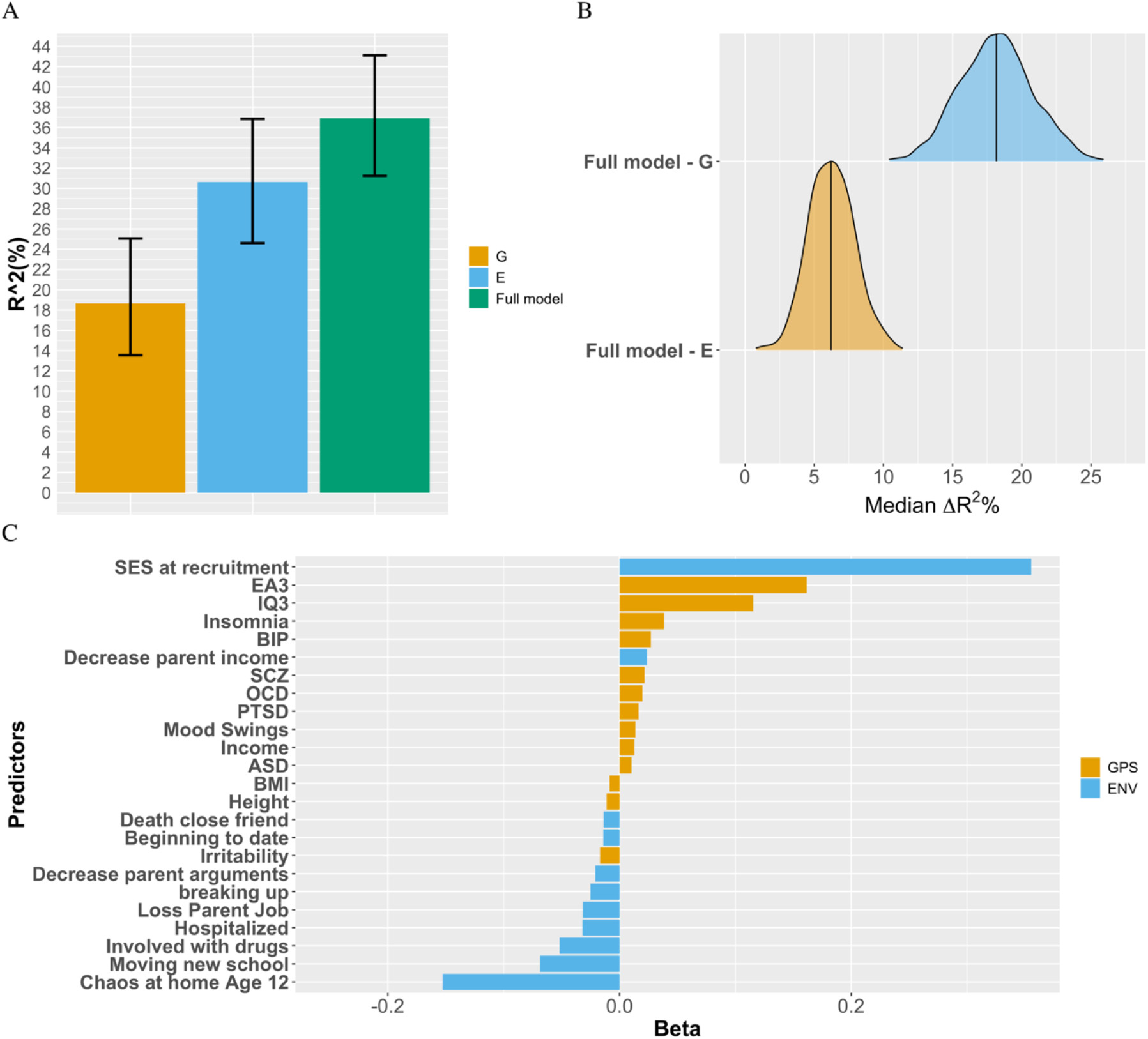
Out of sample prediction of educational achievement. **Panel A** = Hold-out set prediction of EA for the environmental (E), multi-polygenic score (G) and full (joint Environmental and GPS model) prediction models. Error bars are 95% bootstrapped confidence intervals. **Panel B** = bootstrapped median R^2^ difference between the full model and E and G models in the hold-out set, representing independent (non-mediated) genetic effects (Full model - E) and independent environmental effects (Full model - G). **Panel C** = Joint prediction model used in hold-out set prediction (i.e. Full model). Figure shows variables selected via repeated cross-validation in the training set, and relative importance. **Note.** GPS = Genome-wide polygenic score, ENV = Environmental measures. ASD = Autism Spectrum Disorder, BIP = Bipolar Disorder, BMI = Body Mass Index, EA3 = educational attainment, IQ3 = intelligence, OCD = Obsessive Compulsive Disorder, PTSD = Post-Traumatic Stress Disorder, SCZ = Schizophrenia.

### Best-model and coefficient estimation

The best model (full model) selected via repeated cross-validation in the training set included 24 predictors, 13 of which were GPS (orange) while 11 were environments (blue) (Figure 1, panel C). Of these top EA-increasing variables were SES in early life, followed by the GPS for educational attainment (EA3 GPS) and the GPS for intelligence (IQ3 GPS), while the top trait decreasing variable was chaos at home at age 12. In terms of coefficient estimation, partial regression coefficients in post-selection inference analyses (Figure 2 and Supplementary Table S3) showed that EA3 GPS and IQ3 GPS remained significant in the model after adjusting for the other predictors, with the conditional coefficients indicating that 2.7% of the variance in EA was explained by the EA3 GPS (*β* = 0.16; 95% CI = 0.12, 0.20; p = 2.35E-10) and 1.4% by the IQ3 GPS (*β* = 0.12; 95% CI =0.08, 0.15; p = 8.29E-9).

**Figure 2.**
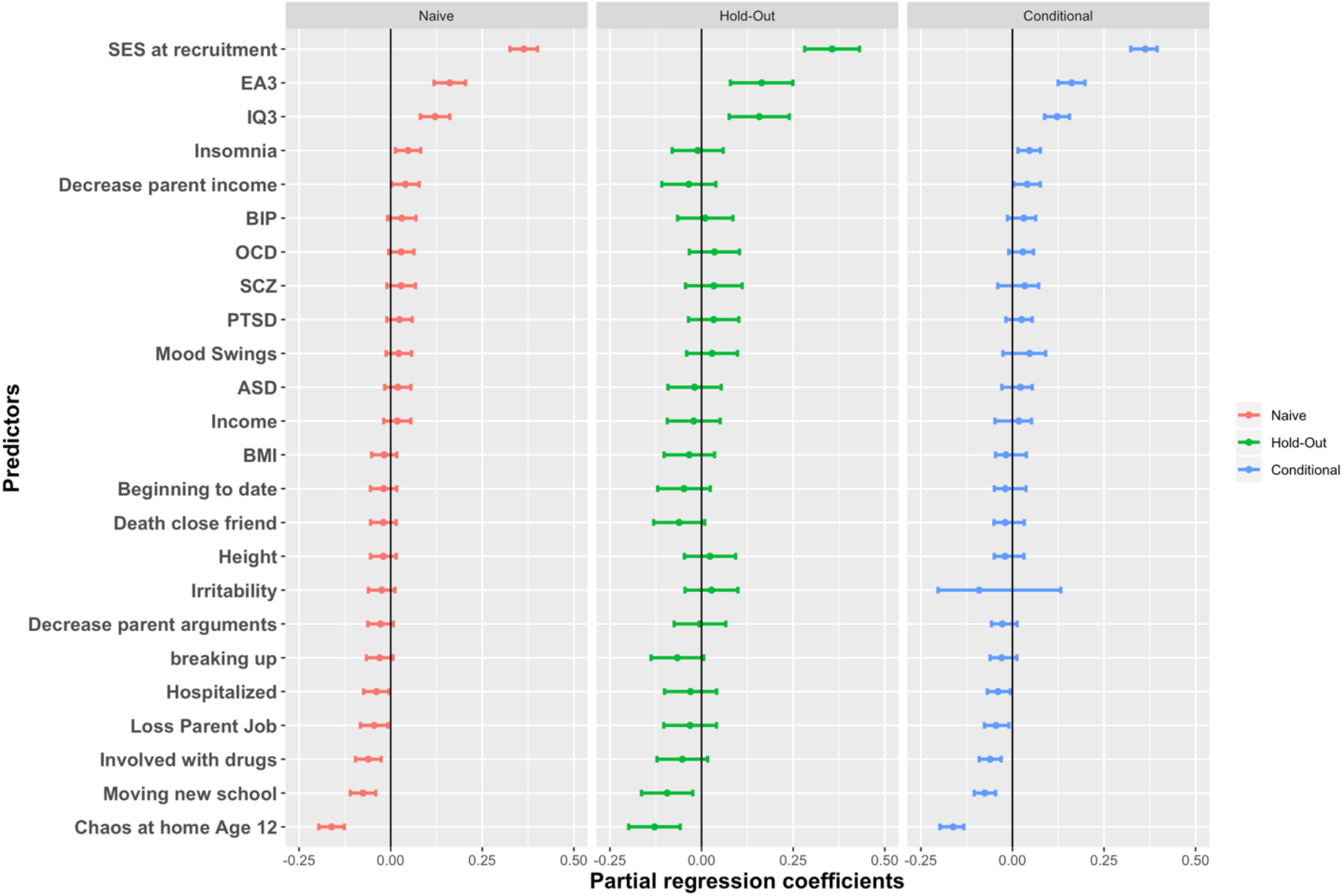
Partial regression coefficients, and 95% CIs around estimates. Naive = partial regression coefficients from multiple regression of selected variables in Training set; Holdout = partial regression coefficients of selected variables in the hold-out set; Conditional = partial regression coefficients of training set for selected variables estimated with a conditional probability from a truncated distribution (see method section).

SES remained the most powerful predictor in the conditional model (*β* = 0.362; 95% CI =0.322, 0.394; p = 1.41E-14). Other environmental exposures that remained significant were ‘chaos at home’ at age 12 (*β* = −0.16; 95% CI =−0.19, −0.13; p = 1.036987e-19) and four life events experienced in the past year (all trait decreasing), including ‘moving to a new school’ (*β* = −0.07; 95% CI −0.10, −0.04; p = 2.22E-5), ‘involved with drugs’ (*β* = −0.06; 95% CI - 0.09, −0.03; p = 7.06E-4), ‘being hospitalized’ (*β* = −0.04; 95% CI −0.07, −0.01; p = 2.77E-2), and ‘loss of a parent job’ (*β* = −0.04; 95% CI −0.08, −0.01; p = 1.97E-02). SES, EA3 GPS, IQ3 GPS and ‘chaos at home’ were significant in all three models (i.e. naive, hold-out and conditional).

### rGE and mediated environmental vs GPS effects

Supplementary Table S2a shows prediction model estimates for all models considered, and Supplementary Table S2b reports nested comparisons of out-of-sample prediction accuracy (R^2^) for the full model vs. E and the full model vs. G. We tested the correlation between the EA predicted values from the G model (G_ea_) and the E model (E_ea_) in the hold-out-set. This was r = .36 (95% CIs = .29, .43), indicating the extent of overlapping information between the G and E models in out-of-sample prediction or, in other words, of rGE (as defined by the variables employed) underlying variation in EA. Then we proceeded to test the extent to which G and E effects on EA were reciprocally mediated (see methods). Supplementary Table S4 shows results of mediation analyses. We found evidence for environmentally mediated genetic effects (indirect path: *β* = 0.16; bootstrapped 95% CI 0.13, 0.20) and genetically mediated environmental effects (indirect path: *β* = 0.10; bootstrapped 95% CI 0.07, 0.13). The effects of G_ea_ on EA (*β* = 0.43; bootstrapped 95% CIs = 0.36, 0.50) were reduced by 38% after introduction of the E_ea_ mediator in the model (*β* = 0.27; bootstrapped 95% CIs = 0.19, 0.34); these effects can be interpreted as the direct G model contributions to EA not accounted for by the E model. In other words, 38% of G effects on EA were explained by environmental mediation. Similarly, the direct E_ea_ effects on EA (*β* = 0.55; bootstrapped 95% CIs = 0.5, 0.60) were subject to a reduction of 18% (*β* = 0.45; bootstrapped 95% CIs = 0.39, 0.51) after introduction of G_ea_ as a mediator in the model, indicating partial genetic mediation of environmental effects (i.e. genetic confounding).

### Discovery of GxE effects and multivariable prediction

We finally tested all possible two-way interactions jointly modelled by means of a hierarchical group lasso procedure (i.e. glinternet, see methods). Out of the possible 528 twoway interactions between all study variables (i.e. interactions between and within sets of GPS and environmental measures), 22 two-way interactions were detected by the hierarchical group-lasso technique (glinternet, Supplementary Table S5), half of which were GxE interactions. Figure 3 depicts an interaction network from the trained glinternet model. Out-of-sample prediction accuracy did not improve over the joint G and E model (R^2^ = 36.5%; 95% CI = 30, 43). We then introduced the 11 GxE interactions found in the full elastic net model to test whether they improved the prediction of EA over the full model that had only considered additive effects of GPS and environmental measures. Similar to the glinternet model, there was no improvement in out-of-sample prediction accuracy (R^2^ = 36.72%; 95% CI = 30.1, 42.83). Supplementary Table S2 shows fit statistics for the glinternet and elastic net models. Supplementary Table S5 reports GxE interactions detected by the hierarchical lasso model.

**Figure 3.**
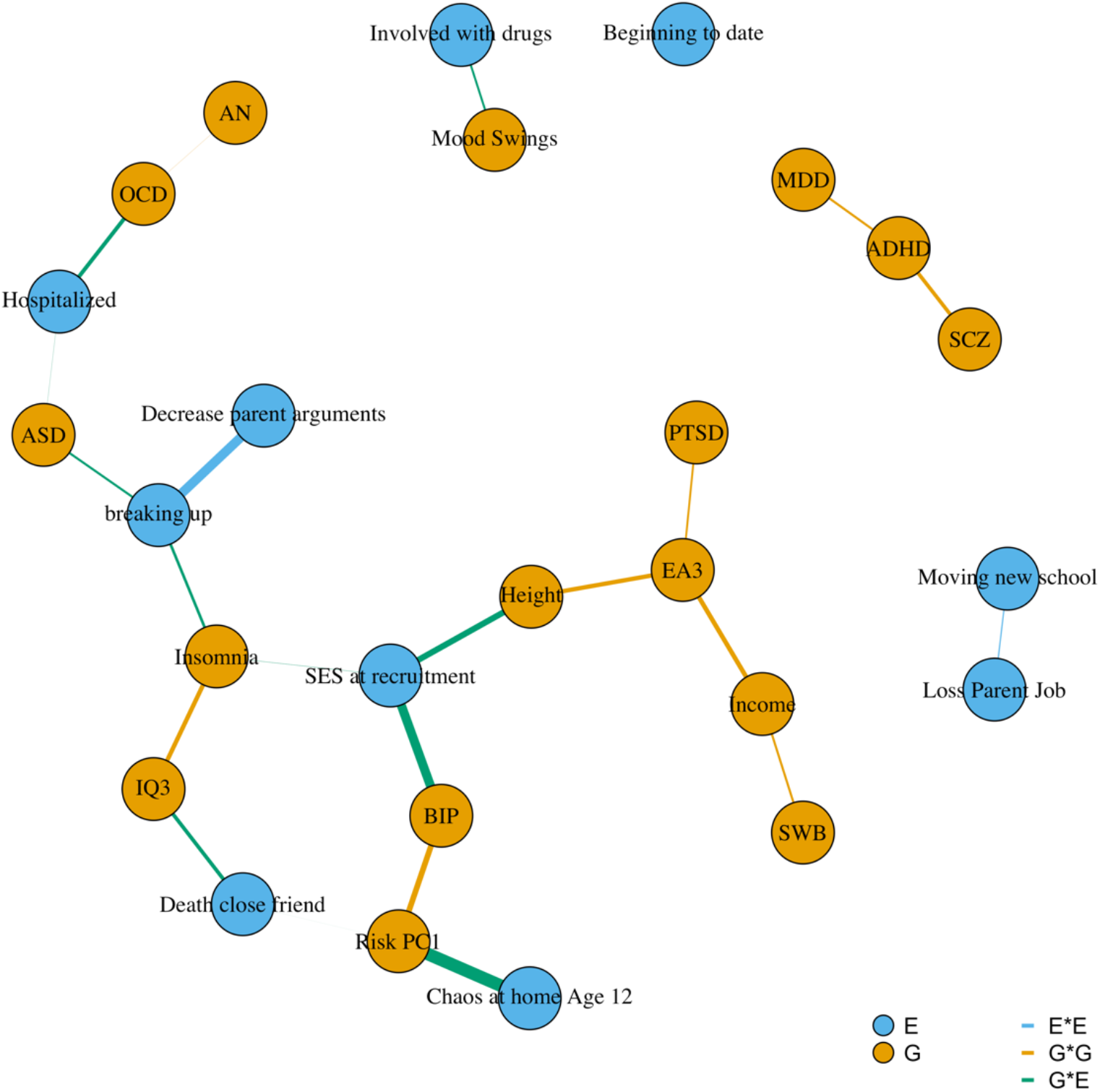
Interaction network of glinternet model. **Note.** E = Environmental measure, G = Genome-wide polygenic score. AN = Anorexia Nervosa, ASD = Autism Spectrum Disorder, ADHD = Attention-Deficit Hyperactivity Disorder, BIP = Bipolar Disorder, EA3 = educational attainment, IQ3 = intelligence, MDD = Major Depressive Disorder, SWB = Subjective Well-Being, OCD = Obsessive Compulsive Disorder, PTSD = Post-Traumatic Stress Disorder, SCZ = Schizophrenia.

## Discussion

We tested the joint prediction accuracy of sets of multiple environmental measures and polygenic scores in prediction models of educational achievement, and considered the effect of their interplay in out-of-sample prediction. Three main findings emerged from our analyses. First, the joint modelling of multiple GPS and related environmental exposures improved the prediction of EA, consistent with theory (39). Second, paralleling quantitative genetic research, we found consistent evidence of rGE effects underlying variation in EA (rGE = .36; 95% CIs = .29, .43). Lastly, we did not find evidence that GxE effects play a role in multivariable prediction of EA.

Our multivariable GPS model alone predicted 18.8% of the variance in EA. Integration of multiple polygenic scores in the same model can be expected to increase as sample size in genome-wide association studies (GWAS) increases (40). Here we constructed GPS in lassosum (41) based on previous observations that lassosum tends to perform better than more traditional approaches (17, 41) for educationally relevant traits. However, other methods for GPS construction can be expected to yield similar results when considering multivariable GPS penalized approaches, with performance of the relative approaches likely to converge as accuracy of GWAS estimates increases.

Of interest were the relative contributions of the single GPS to the model in out-of-sample prediction. In post-selection inference analyses, IQ3 and EA3 were the only GPS independently associated with variation in EA after adjusting for measured environments and polygenic scores. This indicated that both these GPS contributed unique predictive information beyond other related, proxy environmental predictors (e.g. SES, parental educational attainment), and polygenic scores. Similarly, we found that several environments were independently predictive of EA. The best predictor was early life SES, a composite of parental educational attainment, employment status and maternal age at first birth, which alone predicted 13% of the variance in the conditional model. Several life events and chaos at home were also important predictors in the model, with negative independent effects on EA. Polygenic scores, however, improved the prediction of EA on top of the environment with a 19% increase in accuracy (from 30% to 37%). It is noteworthy that EA3 and IQ3 GPS were both significant in post-selection inference models after adjusting for SES, home environment and proximal environmental effects. This suggested that cognitive-relevant GPS independently captured variation beyond environmental variables and variance due to rGE in our model.

A central finding of the current study emerged when we separated direct and indirect effects of the GPS and environmental models by statistically testing for rGE. We found significant G mediation of the prediction of EA by the E model. This is in line with several quantitative and molecular genetics findings (42–44). However, since it would be unreasonable to assume a causal effect of E on G (i.e. E does not change DNA sequence), in the sense employed here G acts as a ‘confounder’ -- in causal modelling parlance, ‘third variable confounding’ -- of E effects on EA (E <-G -> EA). That is, because our G model is associated with both the E model and EA, it partly induces an association between the E model and EA in addition to the independent effects of E on EA. This rGE effect explained 18% of the E effects on EA. Different types of genetic confounding have been described in detail elsewhere (45).

Conversely, we also found evidence of environmental mediation of the G model effects on EA. The E model explained 38% of the GPS model effects on EA. This result is also in line with research in quantitative genetics (46) and molecular genetics (27–29). A growing body of evidence points to the rGE conclusion that genetic effects on cognitive trait variation are partly environmentally mediated (25), which is likely to be due to passive rGE. Passive rGE emerges because parents create a family environment that corresponds to their genotypes and, by extension, also correlates with the genotypes of their offspring. Alternative mechanisms include evocative and active rGE effects. As noted elsewhere (26) these possibilities are not mutually exclusive. However, in order to disentangle these rGE effects, different study designs are needed, for example, looking within families at the effects of maternal and paternal non-transmitted genotypes on child outcomes. Disentangling the different underlying mechanisms to the predicted variance in this regard is an issue for future studies, but out of the scope of the present investigation. Here for the first time we show that reciprocal indirect effects between multivariable E and G prediction models explain a substantial proportion of variation of their direct effects on EA. These results provide converging evidence with recent research looking at rGE underlying parenting and children educational attainment (27). Both genetic confounding and environmental mediation are important factors to take into account in the prediction of EA.

Lastly, we applied a hierarchical group-lasso model (glinternet) to automatically detect twoway interactions. This model helped us to identify GxE effects that show strong hierarchy (see methods), which would have otherwise been difficult to detect due to the great multipletesting burden relative to the sample size of the present study. Furthermore, since glinternet performs shrinkage and grouping before testing for interaction effects, this enabled discovery of interactions that would have been confounded by strong main effects of correlated predictors. In other words, because the coefficients of main effects have been regularized (that is shrunk, see Methods), their fit is reduced, which facilitates the discovery of interaction effects (38). However, neither the full glinternet model including all discovered pairwise interactions, nor the elastic net model including two-way GxE effects, improved out-of-sample prediction accuracy over the full model. One possible explanation for this finding is that GxE effects are typically very small, and that the trade-off between true effect and variance introduced in the model, signal to noise ratio, was too small. It is important to note, however, that application of this method in larger datasets, or using different phenotypes with different genetic architectures, might be fruitful for hypothesis-free GxE discovery.

Our findings parallel those from quantitative genetic studies in humans, which reported widespread rGE, but little evidence for GxE. Even quantitative genetic studies of nonhuman animals found little evidence for GxE (47), even though they afford more powerful research designs for testing interactions. One possible explanation is that measured GxE effects are unsystematic and idiosyncratic. In other words, nonlinear effects might be too noisy in multivariable prediction models such as those employed here, suggesting that GxE might be more important to be considered for inference in statistical models, rather than in the context of predicting phenotypic differences. In other words, statistically modelling of GxE interactions might be a powerful tool for explanatory modelling (theory building or testing), for which effect estimates do not need to be sizeable to be informative, while for prediction modelling the opposite is true. As large multidimensional biobank datasets become increasingly available, the integration of multi-omics data with multiple environmental measures will become more common. Here, we provide an indication of the effects of integrating multiple GPS and environmental measures in prediction models of EA and the effect that their interplay has on prediction accuracy in a population cohort of adolescents aged 16 years.

Our results are limited to the variables employed in our analyses. For example, we modelled exposures that are typically defined as environmental; however, many other variables can be argued to capture environmental influences. Likewise, we included a broad range of GPS that are currently available as the most predictive for cognitive, psychiatric and anthropometric outcomes, but other GPS may also be relevant. Finally, we focused here on EA but predictive models of other complex traits are likely to yield different results, because EA shows comparatively great shared environmental influences (30). This suggests that rGE is likely to be stronger for EA than for other behavioural traits, such as personality traits and social-emotional competencies. Regarding our analytical approach, we focused on GxE interactions that obeyed strong hierarchy as identified by the group lasso technique. Future studies could relax this assumption and include interactions where one of the main effect sizes is not significant. Finally, although it is a strength of our study that we used measured environmental exposures, we note that methods for inferring GxE without measured environmental data are emerging that have reported GxE for some complex traits (48). The extent to which these effects are systematic, stable, and generalizable to EA remains to be determined.

In conclusion, we found evidence for rGE in prediction models of EA that systematically tested the interplay between polygenic scores and measured environments within a hypothesis-free multivariable prediction framework. When integrating multiple GPS and environmental measures into comprehensive prediction models, their correlations must be taken into account. Separate effects of environmental and polygenic scores cannot just be assumed to add up because pervasive rGE affects prediction.

## Methods

### Sample

We test our models using data from 16 year olds from the UK Twin Early Development Study (TEDS; 49), a large longitudinal study involving 16,810 pairs of twins born in England and Wales between 1994-1996, with DNA data available for 10,346 individuals (including 3,320 dizygotic twin pairs and 7,026 unrelated individuals). Ethical approval for TEDS has been provided by the King’s College London Ethics Committee (reference: PNM/09/10– 104). Parental consent was obtained before data collection. Genotypes for the 10,346 individuals were processed with stringent quality control procedures followed by SNP imputation using the Haplotype Reference Consortium (release 1.1) reference panels. Current analyses were limited to the genotyped and imputed sample of 7,026 unrelated individuals. Following imputation, we excluded variants with minor allele frequency <0.5%, Hardy-Weinberg equilibrium p-values of <1 × 10-5. To ease computational demands, we selected variants with an info score of 1, resulting in 515,000 SNPs used for analysis (see the supplementary information for a full description of quality control and imputation procedures).

### Measures

#### Dependent measure: Educational Achievement

Educational achievement was measured as the self-reported mean grade of three core subjects (English, math and science) scored by the individuals at age 16 in their standardized UK General Certificate of Secondary Education (GCSE) exams.

EA was operationalized as the mean grade of the three compulsory subjects, with results coded from 4 (G, or lowest grade) to 11 (A+, or highest grade). These self-report measures are highly replicable and show high genetic and phenotypic correlations with teacher scores(50). The variable distribution was slightly left skewed (similar to the national average) and subject to a rank based inverse normal transformation to approximate a normal distribution.

#### Environmental measures

##### Socio economic status

SES at recruitment (mean age = 18 months) was calculated as a composite of mother and father qualification levels ranging from 1 = ‘no qualifications’ to 8 = ‘postgraduate qualification’, mother and father employment status (51), and mother’s age on birth of first child.

##### Chaos at home

as a measure of home environment a shortened version of the Confusion, Hubbub and Order Scale (52) was used to measure children’s perception of chaos in the family environment at age 12. Children rated the extent to which they agree (range: ‘not true’, ‘quite true’ or ‘very true’) to six items: ‘I have a regular bedtime routine’ (reversed coded), ‘You can’t hear yourself think in our home’, ‘It’s a real zoo in our home’, ‘We are usually able to stay on top of things’ (reversed coded), ‘There is usually a television turned on somewhere in our home’ and ‘The atmosphere in our house is calm’ (reverse coded). The Chaos score was computed as the mean of the rated items.

##### Life events

Self-reported life events experienced in the past year were measured (at age 16) using a shortened version of the Coddington life events (53). Individuals had to report on 20 items that might have happened in the past year, by responding yes (coded as 1) if the event had happened or no (coded as 0) if it didn’t happen. Items included stochastic, proximal events such as “death of a close friend or relative”, “being hospitalized”, as well as family-wide events e.g. “loss of a parent job”, “decrease in parental income”. When considering prediction of educational achievement, educationally relevant items were removed from the models (i.e. “failing exam” and “outstanding achievement”). Items being endorsed by fewer than 100 people were discarded from analyses. A total of 11 life events were retained in analyses. All items were considered separately in prediction models (i.e. they were not aggregated in a scale).

Supplementary Table S1 reports descriptive statistics for variables employed in this study, separately by training and test sets.

##### Genome-wide polygenic scores (GPS)

GPS for 20 cognitive, anthropometric and psychopathological traits were constructed using Lassosum (41). Lassosum is a penalized regression approach applied to GWAS summary statistics. In lassosum we try to minimize the following loss function:

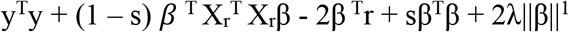

Here *λ* controls the L1 penalty (L1 norm, (54)). The notation ||β||^1^ describes the L1 norm of a coefficient vector β, defined as

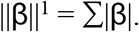

While s is another tuning parameter controlling the L2 penalty (|β|^2^, the sum of the squared betas). Here *s* has the additional constraint of being between 0 and 1. When *λ* = 0 and s = 1 the problem becomes unconstrained.

Tuning parameters, *λ* and s, are chosen in the validation step (this is akin to optimization that can be performed in p-value thresholding methods). We used our training set to perform parameter tuning optimizing (with respect to R^2^) polygenic scores against EA. LD was accounted for via a reference panel, here the same as the training set sample, and estimation of LD blocks was performed using LD regions defined in (55).

We created cognitive and educationally relevant polygenic scores for educational attainment (16), intelligence (56), and income (57). We also created polygenic scores for mental health-related traits: autism spectrum disorder (58), major depressive disorder (MDD; 59), bipolar disorder (BIP; 60), schizophrenia (SCZ; 61), attention deficit hyperactivity disorder (ADHD; 62), obsessive compulsive disorder (OCD; 63), anorexia nervosa (AN; 64), post-traumatic stress disorder (PTSD; 65), depressive symptoms (66), mood swings (67), subjective wellbeing (66), neuroticism (68), irritability (67), insomnia (69), and risk taking (70). Finally, we created polygenic scores for height and BMI (71). Supplementary Table S6 reports information on these summary statistics, while Supplementary Table S7 reports parameter tunings for the lassosum GPS.

### Analyses

All variables were first regressed on age, sex, 10 genetic principal components and genotyping chip. The obtained standardized residuals were used in all subsequent analyses.

#### Penalised regression

We fit elastic net regularization (72) models for EA. Elastic Net minimizes the residual sum of squares (RSS) subject to the L1 penalty, which consist of the sum of the absolute coefficients and the L2 penalty which consist of the sum of the squared coefficients, performing parameters shrinkage and variable selection at the same time (72).

Elastic net tries to minimise the following loss function:

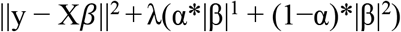

where

||y - X*β*||^2^ is the residual sum of squares
|β|^2^ is the sum of the squared betas (the L2 penalty)
|β|^1^ is the sum of the absolute betas (the L1 penalty)

Here α determines the mixing of penalties, where the first parameter introduces sparsity while the second shrinks correlated predictors towards each other. *λ* is a tuning parameter that control the effect of the penalty terms over the regression coefficients. When α = 1 the solution is equivalent to a LASSO regression, while when α = 0 the solutions is equivalent to a Ridge regression.

For every alpha multiple Lambdas exists, and the optimal combination of tuning parameters is determined by repeated cross-validation. For every model tested we split the sample in a training test (80%) and a hold-out set (20%) treated independently with respect to data wrangling. In the training set we perform 10-fold repeated cross-validation to select the model that minimises the Root Mean Square Error (RMSE) – that is the tuning parameter for which the cross-validation error is the smallest. The model performance is then assessed by the variance explained (R^2^) in the independent hold-out test set. After separately training penalized regression models we compare the best E vs G vs joint prediction models using standard prediction accuracy and loss function indices in the hold-out set.

##### Bootstrapping

For every model tested we sampled with replacement from the data (1000 times) to calculate bootstrapped confidence intervals for the out-of-sample prediction accuracy (R^2^). Confidence intervals were defined as the 2.5th and 97.5th percentiles of the R^2^ distributions obtained from bootstrapping. Then we calculated median differences for each pairwise R^2^ distribution between models, and calculated 95% confidence levels as the 2.5 and 97.5 percentiles of these distributions. If the interval didn’t contain 0 we concluded that the pairwise model R^2^ difference was significantly different from 0 with a α level of .05.

##### Post selection inference

For every model tested we conducted inference of models coefficients after selection of most informative predictors performed by Elastic Net, that is effect sizes, p-values and confidence levels around the prediction estimates.

Post-selection inference (37) refers to the practice of attempting to accurately estimate prediction coefficients after a model selection has been performed. If we fit the optimal model’s selected predictors in a multiple regression model in the training set (that is where the selection has been performed) our confidence in the estimates will tend to be over-optimistic. On the other hand, estimation of these parameters in a hold-out set would not be subject to this problem. The hold-out set, however, will typically be smaller than the training set, leading to wide confidence intervals. In addition, the results will be dependent on the random split (80-20) performed. A third way is to calculate P-values conditional to the selection that has been made in the training set.

Briefly, after selection is performed, accurate estimation of a given partial regression coefficient can be approximated by a truncated normal distribution:

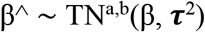

With mean β, variance **τ**^2^ and boundaries of the truncated normal distribution (TN) ‘a’ and ‘b’ given by the data and the selection procedure, in this case the predictors, the active set (the variables with non 0 coefficients selected by our model) and *λ* (37). We refer elsewhere to a thorough discussion of the topic (73), with a focus on lasso like approaches. Here we compare results from the three procedures: the ‘naive’ estimation of partial regression coefficients in the training set, estimation of coefficients in the hold-out set, and the conditional estimation of p-values performed using the R package ‘SelecitveInference’ (74).

#### rGE

We quantified rGE in two ways. First, the hold-out set predicted EA values from the GPS (enceforth G_ea_) and environmental (henceforth E_ea_) models can be tested for correlation. In this sense the covariance between these variables would be an indication of overlapping information between E and G underlying EA, i.e. r_G,E_ = cor(G_ea_,E_ea_). Second, another way to quantify rGE is by modelling E and G effects in a mediation model (Figure 4), considering the indirect effects of G on EA via E, and vice versa the indirect effects of E on EA via G. We used the predicted EA values from the GPS and environmental models (i.e. G_ea_ and E_ea_) to test mediation models in the hold-out set. We fit a structural equation model (SEM) in ‘lavaan’ (71) to test whether and to what extent E and G effects on EA were reciprocally mediated. Panel B is a schematic representation of a mediation model, where βC is the effect of a predictor X on an outcome Y, βa the effect of X on the mediator (M), and βb the effect of M on Y after adjusting for X. βC’ corresponds to the effects of the predictor on the outcome when controlling for the mediator (i.e. when the full equation is estimated). If the effects are reduced (partial mediation) or are not different from 0 (full mediation) then there is evidence for mediation. We quantify the proportion of the mediated effects as (β C- βC’) / βC (72) and test for significance of the indirect path using bootstrapping (with 1000 repetitions).

**Figure 4.**
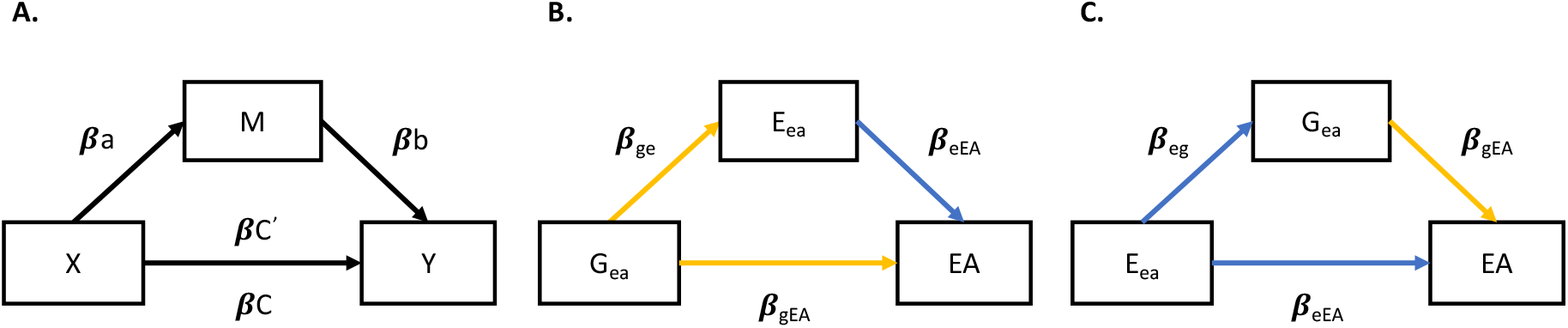
*Panel A* = schematic representation of mediation analysis; *βC* = effect of a predictor X on an outcome Y; *βa* = effect of X on a mediator (M); *βb* = effect of M on Y after adjusting for X; *βC’* = effect of X on Y after adjusting for M. *Panel B* = Directed acyclic graph (DAG) showing E_ea_ mediated effects of G_ea_ on EA in the hold out-set; *β*_ge_ = causal path between G_ea_ and E_ea_ equivalent to r_G,E_; *β*_eEA_ = direct independent E_ea_ effects on EA; *β_eEA_* = total G_ea_ effects on EA. *Panel C* = DAG showing G_ea_ mediated effects on EA (genetic confounding, see methods and discussion); *β*_eg_ = causal path between E_ea_ and G_ea_ equivalent to r_G,E_; *β*_gEA_ = direct independent G_ea_ effects on EA; *β_eEA_* = total E_ea_ effects on EA. **Note.** Yellow paths represent G model effects, blue paths represent E model Effects.

Figure 4 represent direct and indirect effects of the G model effects on EA mediated by E (panel B), and of the E model effects on EA mediated by G (panel C). While panel B represents a causal model where we estimate the environmentally mediated G effects on EA, panel C is a statistical abstraction since it would be unreasonable to assume a causal relationship of E on G. Here we model G as mediator to estimate the third variable confounding effects underlying the relationship between E and EA, as mediating and confounding effects have been shown to be equivalent in a linear context (75).

#### GxE

After fitting the joint GPS and environmental models, we apply a hierarchical lasso procedure to automatically search the feature space for interactions, and retrain our models introducing GxE interactions. With 33 predictors there is a total of 33(33-1)/2 = 528 possible 2-ways interactions. Testing all models separately would imply a multiple testing burden (e.g. bonferroni correction .05/820 = 9E-5), in addition to the expected low signal to noise ratio for GxE effects. Here we employ a hierarchical group lasso approach to automatically search for two-way interactions, implemented in the R package ‘glinternet’ (38) (group-lasso interaction network). Glinternet leverages group lasso, an extension of LASSO, to perform variable selection on groups of variables, dropping or retaining them in the model at the same time, to select interactions. As noted above, the L1 regularization produces sparsity. Glinternet uses a group lasso for the variables and variable interactions, which introduces a strong hierarchy: an interaction between two variables can only be picked by the model if both variables are also selected as main effects. That is, interactions between two predictors are not considered unless both predictors have non-zero coefficients in the model. Once two-way interactions obeying strong hierarchy were identified, we selected GxE interactions (i.e. GPS that interact with environmental variables) and reintroduced them in our best elastic net models to test whether the test set prediction accuracy improved beyond the full (joint E-G) prediction model.

## Acknowledgements

We gratefully acknowledge the ongoing contribution of the participants in the Twins Early Development Study (TEDS) and their families. TEDS is supported by a programme grant to RP from the UK Medical Research Council (MR/M021475/1 and previously G0901245), with additional support from the US National Institutes of Health (AG046938). The research leading to these results has also received funding from the European Research Council under the European Union’s Seventh Framework Programme (FP7/2007-2013)/grant agreement n° 602768 and ERC grant agreement n° 295366. RP is supported by a Medical Research Council Professorship award (G19/2). This project has received funding from the European Union’s Horizon 2020 research and innovation programme under the Marie Sklodowska-Curie grant agreement no. 721567. JRIC is supported in part by the UK National Institute for Health Research (NIHR) as part of the Maudsley Biomedical Research Centre (BRC). This study represents independent research partly funded by the NIHR BRC at South London and Maudsley NHS Foundation Trust and King’s College London. The views expressed are those of the author(s) and not necessarily those of the NHS, the NIHR or the Department of Health and Social Care. SvS is supported by a Jacobs Fellowship (2017-2019).

High performance computing facilities were funded with capital equipment grants from the GSTT Charity (TR130505) and Maudsley Charity (980).

## References

1. K. Asbury, R. Plomin (2013) G is for genes: what genetics can teach us about how we teach our children. (Wiley, Oxford).

2. K. Rimfeld et al., The stability of educational achievement across school years is largely explained by genetic factors. NPJ science of learning 3, 16 (2018).

3. T. J. C. Polderman et al., Meta-analysis of the heritability of human traits based on fifty years of twin studies. Nature genetics 47, 702 (2015).

4. R. Plomin, C. S. Bergeman, The nature of nurture: Genetic influence on “environmental” measures. Behavioral and Brain Sciences 14, 373–386 (1991).

5. R. Plomin, J. C. DeFries, J. C. Loehlin, Genotype–environment interaction and correlation in the analysis of human behavior. Psychological bulletin 84, 309 (1977).

6. D. W. Belsky et al., Genetic analysis of social-class mobility in five longitudinal studies. Proceedings of the National Academy of Sciences 115, E7275–E7284 (2018).

7. S. Selzam et al., Evidence for gene-environment correlation in child feeding: Links between common genetic variation for BMI in children and parental feeding practices. PLoS genetics 14, e1007757 (2018).

8. A. Abdellaoui et al., Genetic correlates of social stratification in Great Britain. Nature human behaviour, 1–21 (2019).

9. J. Y. Lau, T. C. Eley, Disentangling gene-environment correlations and interactions on adolescent depressive symptoms. Journal of Child Psychology and Psychiatry 49, 142–150 (2008).

10. A. Caspi et al., Influence of life stress on depression: moderation by a polymorphism in the 5-HTT gene. Science 301, 386–389 (2003).

11. R. Border et al., No support for historical candidate gene or candidate gene-by-interaction hypotheses for major depression across multiple large samples. American Journal of Psychiatry 176, 376–387 (2019).

12. D. M. Dick et al., Candidate gene-environment interaction research: Reflections and recommendations. Perspectives on Psychological Science 10, 37–59 (2015).

13. L. E. Duncan, M. C. Keller, A critical review of the first 10 years of candidate gene-by-environment interaction research in psychiatry. American Journal of Psychiatry 168, 1041–1049 (2011).

14. P. M. Visscher et al., 10 years of GWAS discovery: biology, function, and translation. The American Journal of Human Genetics 101, 5–22 (2017).

15. N. R. Wray et al., Research review: polygenic methods and their application to psychiatric traits. Journal of Child Psychology and Psychiatry 55, 1068–1087 (2014).

16. J. J. Lee et al., Gene discovery and polygenic prediction from a genome-wide association study of educational attainment in 1.1 million individuals. Nature genetics 10.1038/s41588-018-0147-3 (2018).

17. A. Allegrini et al., Genomic prediction of cognitive traits in childhood and adolescence. Molecular psychiatry 24, 819 (2019).

18. N. Mullins et al., Polygenic interactions with environmental adversity in the aetiology of major depressive disorder. Psychological medicine 46, 759–770 (2016).

19. P. B. Barr et al., Polygenic Risk for Alcohol Misuse is Moderated by Romantic Partnerships. Addiction (Abingdon, England) (2019).

20. S. H. Barcellos, L. S. Carvalho, P. Turley, Education can reduce health differences related to genetic risk of obesity. Proceedings of the National Academy of Sciences 115, E9765–E9772 (2018).

21. J. A. Pasman, K. J. Verweij, J. M. Vink, Systematic Review of Polygenic Gene– Environment Interaction in Tobacco, Alcohol, and Cannabis Use. Behavior genetics, 1–17 (2019).

22. J. R. Coleman, E. Krapohl, T. C. Eley, G. Breen, Individual and shared effects of social environment and polygenic risk scores on adolescent body mass index. Scientific reports 8, 6344 (2018).

23. W. J. Peyrot et al., Does childhood trauma moderate polygenic risk for depression? A meta-analysis of 5765 subjects from the psychiatric genomics consortium. Biological psychiatry 84, 138–147 (2018).

24. H. Dobewall et al., Gene–environment correlations in parental emotional warmth and intolerance: genome-wide analysis over two generations of the Young Finns Study. Journal of Child Psychology and Psychiatry 60, 277–285 (2019).

25. D. W. Belsky et al., Genetic analysis of social-class mobility in five longitudinal studies. Proceedings of the National Academy of Sciences 115, E7275–E7284 (2018).

26. E. Krapohl et al., Widespread covariation of early environmental exposures and trait-associated polygenic variation. Proceedings of the National Academy of Sciences 114, 11727–11732 (2017).

27. J. Wertz et al., Genetics of nurture: A test of the hypothesis that parents’ genetics predict their observed caregiving. Developmental psychology (2019).

28. T. C. Bates et al., The nature of nurture: Using a virtual-parent design to test parenting effects on children’s educational attainment in genotyped families. Twin Research and Human Genetics 21, 73–83 (2018).

29. A. Kong et al., The nature of nurture: Effects of parental genotypes. Science (New York, N.Y.) 359, 424–428 (2018).

30. S. Selzam et al., Comparing within- and between-family polygenic score prediction. BioRxiv, 605006 (2019).

31. R. Cheesman et al., Comparison of adopted and non-adopted individuals reveals gene-environment interplay for education in the UK Biobank. bioRxiv, 707695 (2019).

32. E. Krapohl et al., Multi-polygenic score approach to trait prediction. Molecular psychiatry 23, 1368–1374 (2018).

33. E. Krapohl et al., The high heritability of educational achievement reflects many genetically influenced traits, not just intelligence. Proceedings of the National Academy of Sciences 111, 15273–15278 (2014).

34. K. Rimfeld, Y. Kovas, P. S. Dale, R. Plomin, True grit and genetics: Predicting academic achievement from personality. Journal of personality and social psychology 111, 780 (2016).

35. E. Krapohl et al., Phenome-wide analysis of genome-wide polygenic scores. Molecular psychiatry 21, 1188 (2015).

36. J. Friedman, T. Hastie, R. Tibshirani, Regularization paths for generalized linear models via coordinate descent. Journal of statistical software 33, 1 (2010).

37. J. Taylor, R. J. Tibshirani, Statistical learning and selective inference. Proceedings of the National Academy of Sciences 112, 7629–7634 (2015).

38. M. Lim, T. Hastie, Learning interactions via hierarchical group-lasso regularization. Journal of Computational and Graphical Statistics 24, 627–654 (2015).

39. F. Dudbridge, N. Pashayan, J. Yang, Predictive accuracy of combined genetic and environmental risk scores. Genetic epidemiology 42, 4–19 (2018).

40. F. Dudbridge, Power and predictive accuracy of polygenic risk scores. PLoS genetics 9, e1003348 (2013).

41. T. S. H. Mak, R. M. Porsch, S. W. Choi, X. Zhou, P. C. Sham, Polygenic scores via penalized regression on summary statistics. Genetic epidemiology 41, 469–480 (2017).

42. R. Plomin, J. C. DeFries, V. S. Knopik, J. M. Neiderhiser, Top 10 replicated findings from behavioral genetics. Perspectives on psychological science 11, 3–23 (2016).

43. R. Plomin, Genotype-environment correlation in the era of DNA. Behavior genetics 44, 629–638 (2014).

44. E. Krapohl, R. Plomin, Genetic link between family socioeconomic status and children’s educational achievement estimated from genome-wide SNPs. Molecular psychiatry 21, 437 (2015).

45. J.-B. Pingault et al., Estimating the sensitivity of associations between risk factors and outcomes to shared genetic effects. bioRxiv, 592352 (2019).

46. R. Plomin, C. S. Bergeman, The nature of nurture: Genetic influence on “environmental” measures. Behavioral and Brain Sciences 14, 373–386 (2011).

47. V. S. Knopik, J. M. Neiderhiser, J. C. DeFries, R. Plomin, Behavioral genetics (Macmillan Higher Education, 2016).

48. J. Sulc et al., Maximum likelihood method quantifies the overall contribution of gene-environment interaction to complex traits: an application to obesity traits. bioRxiv, 632380 (2019).

49. K. Rimfeld et al., Twins Early Development Study: A Genetically Sensitive Investigation into Behavioral and Cognitive Development from Infancy to Emerging Adulthood. Twin Research and Human Genetics, 1–6 (2019).

50. K. Rimfeld et al., Teacher assessments during compulsory education are as reliable, stable and heritable as standardized test scores. Journal of Child Psychology and Psychiatry (2019).

51. O. o. P. a. C. Surveys, Standard occupational classification. Her Majesty’s Stationery Office volume 3 (1991).

52. A. P. Matheny Jr, T. D. Wachs, J. L. Ludwig, K. Phillips, Bringing order out of chaos: Psychometric characteristics of the confusion, hubbub, and order scale. Journal of Applied Developmental Psychology 16, 429–444 (1995).

53. R. D. Coddington, The significance of life events as etiologic factors in the diseases of children: II. A study of a normal population. Journal of psychosomatic research (1972).

54. R. Tibshirani, Regression shrinkage and selection via the Lasso. J R Stat Soc Ser B Methodol, 267–288 (1996).

55. T. Berisa, J. K. Pickrell, Approximately independent linkage disequilibrium blocks in human populations. Bioinformatics 32, 283–285 (2016).

56. J. E. Savage et al., Genome-wide association meta-analysis in 269,867 individuals identifies new genetic and functional links to intelligence. Nature genetics 50, 912-919 (2018).

57. W D. Hill et al., Molecular Genetic Contributions to Social Deprivation and Household Income in UK Biobank. Current Biology 26, 3083–3089 (2016).

58. J. Grove et al., Identification of common genetic risk variants for autism spectrum disorder. Nature genetics 51, 431 (2019).

59. N. R. Wray et al., Genome-wide association analyses identify 44 risk variants and refine the genetic architecture of major depression. Nature genetics 50, 668 (2018).

60. E. A. Stahl et al., Genome-wide association study identifies 30 loci associated with bipolar disorder. Nature genetics 51, 793 (2019).

61. A. F. Pardiñas et al., Common schizophrenia alleles are enriched in mutation-intolerant genes and in regions under strong background selection. Nature genetics 50, 381 (2018).

62. D. Demontis et al., Discovery of the first genome-wide significant risk loci for attention deficit/hyperactivity disorder. Nature genetics 51, 63 (2019).

63. I. O. C. D. F. Genetics et al., Revealing the complex genetic architecture of obsessive–compulsive disorder using meta-analysis. Molecular psychiatry 23, 1181 (2018).

64. L. Duncan et al., Significant locus and metabolic genetic correlations revealed in genome-wide association study of anorexia nervosa. American journal of psychiatry 174, 850–858 (2017).

65. L. E. Duncan et al., Largest GWAS of PTSD (N= 20 070) yields genetic overlap with schizophrenia and sex differences in heritability. Molecular psychiatry 23, 666 (2018).

66. A. Okbay et al., Genetic variants associated with subjective well-being, depressive symptoms, and neuroticism identified through genome-wide analyses. Nature genetics 48, 624 (2016).

67. C. Seed (2017) Hail: An Open-Source Framework for Scalable Genetic Data.

68. M. Luciano et al., Association analysis in over 329,000 individuals identifies 116 independent variants influencing neuroticism. Nature genetics 50, 6 (2018).

69. P. R. Jansen et al., Genome-wide analysis of insomnia in 1,331,010 individuals identifies new risk loci and functional pathways. Nature genetics 51, 394 (2019).

70. R. K. Linnér et al., Genome-wide association analyses of risk tolerance and risky behaviors in over 1 million individuals identify hundreds of loci and shared genetic influences. Nature genetics 51, 245 (2019).

71. L. Yengo et al., Meta-analysis of genome-wide association studies for height and body mass index in~ 700000 individuals of European ancestry. Human molecular genetics 27, 3641–3649 (2018).

72. H. Zou, T. Hastie, Regularization and variable selection via the elastic net. Journal of the royal statistical society: series B (statistical methodology) 67, 301–320 (2005).

73. K. Liu, J. Markovic, R. Tibshirani, More powerful post-selection inference, with application to the lasso. arXiv preprint arXiv:1801.09037 (2018).

74. J. R. Loftus, Selective inference after cross-validation. arXiv preprint arXiv:1511.08866 (2015).

75. D. P. MacKinnon, J. L. Krull, C. M. Lockwood, Equivalence of the mediation, confounding and suppression effect. Prevention science 1, 173–181 (2000).

