## Supplementary Information for "Multivariable G-E interplay in the prediction of educational achievement"

**Supplementary figures**

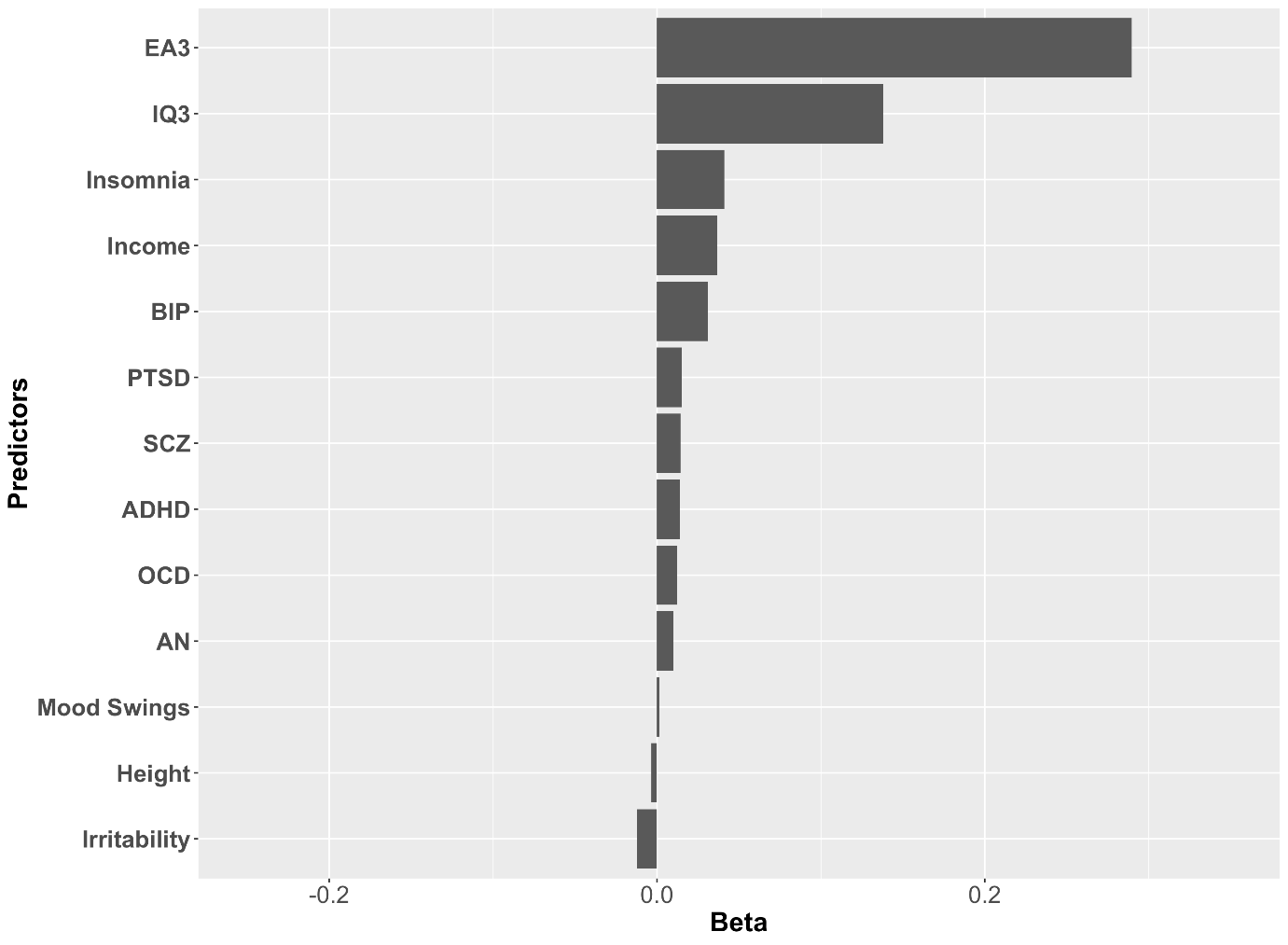

**Supplementary figure 1.** G model used in hold-out set prediction. Figure shows variables importance for the best G model selected via repeated cross-validation in the training set.

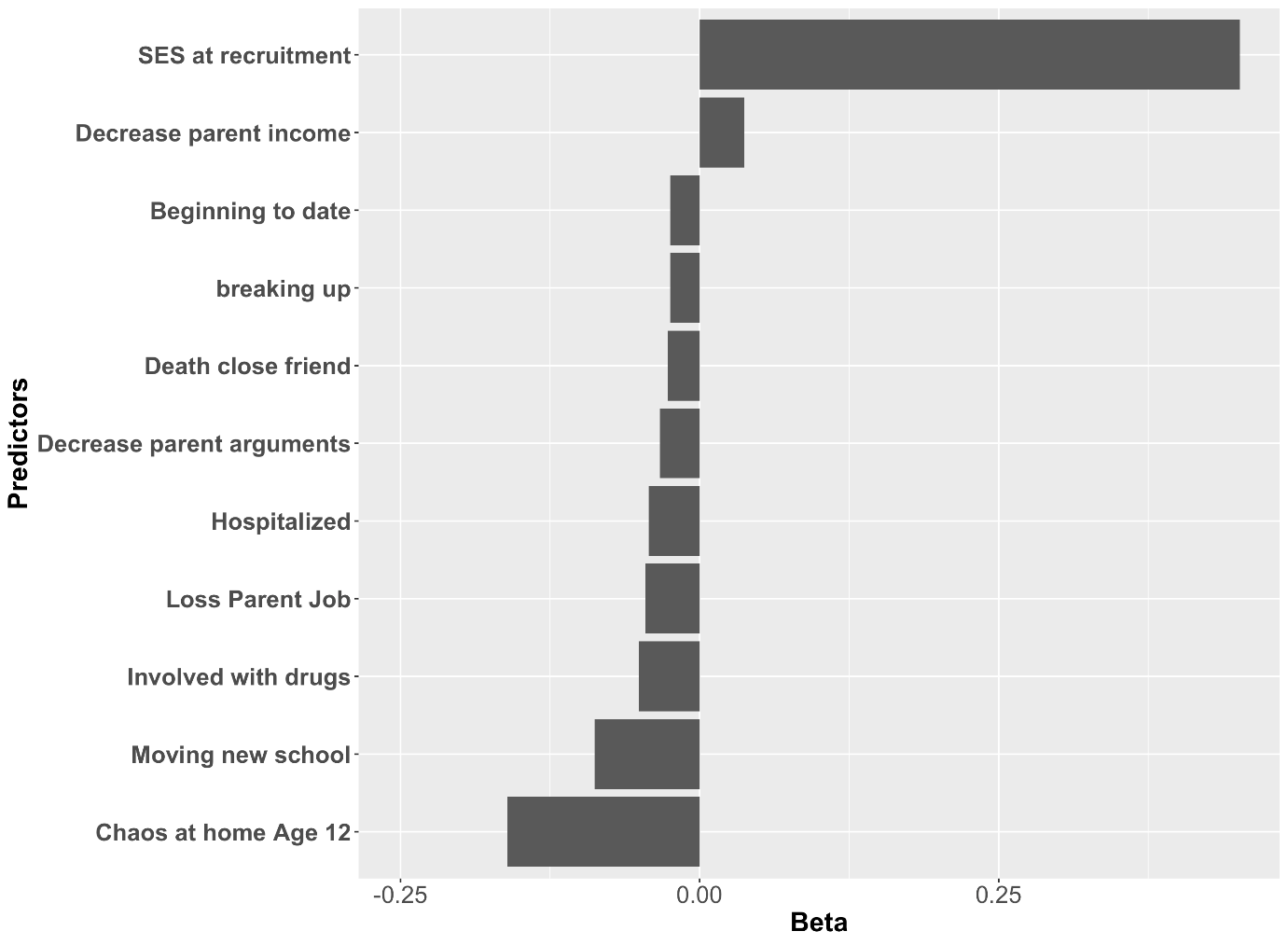

**Supplementary figure 2.** E model used in hold-out set prediction. Figure shows variables importance for the best E model selected via repeated cross-validation in the training set.

**Supplementary information**

**QC and genotyping protocol**

DNA for 8,743 individuals (including 3,722 dizygotic co-twin samples) was extracted from saliva and buccal cheek swab samples and hybridized to HumanOmniExpressExome-8v1.2 genotyping arrays at the Institute of Psychiatry, Psychology and Neuroscience Genomics & Biomarker Core Facility, London, UK. The raw image data from the array were normalized, pre-processed, and filtered in GenomeStudio according to Illumina Exome Chip SOP v1.4. (<http://confluence.brc.iop.kcl.ac.uk:8090/display/PUB/Production+Version%3A+Illumina+Exome+Chip+SOP+v1.4>). In addition, prior to genotype calling, 919 multi-mapping SNPs and 501 samples with callrate <0.95 were removed. The ZCALL program was used to augment the genotype calling for samples and SNPs that passed the initial QC.

DNA from 3,747 samples was extracted from buccal cheek swabs and genotyped at Affymetrix, Santa Clara, California, USA. From this sample, 3,665 samples were successfully hybridized to AffymetrixGeneChip 6.0 SNP genotyping arrays (<http://www.affymetrix.com/support/technical/datasheets/genomewide_snp6_datasheet.pdf>) using experimental protocols recommended by the manufacturer (Affymetrix Inc., Santa Clara, CA). The raw image data from the arrays were normalized and pre-processed at the Wellcome Trust Sanger Institute, Hinxton, UK for genotyping as part of the Wellcome Trust Case Control Consortium 2 (<https://www.wtccc.org.uk/ccc2/>) according to the manufacturer’s guidelines (http://www.affymetrix.com/support/downloads/manuals/genomewidesnp6_manual.pdf). Genotypes for the Affymetrix arrays were called using CHIAMO (https://mathgen.stats.ox.ac.uk/genetics_software/chiamo/chiamo.html).

After initial quality control and genotype calling, the same quality control was performed on the samples genotyped on the Illumina and Affymetrix platforms separately using PLINK (Chang et al., 2015; Purcell et al., 2007), R (R Core Team, n.d.), BCFtools (Li, 2011), and EIGENSOFT (Patterson, Price, & Reich, 2006; Price et al., 2006).

Samples were removed from subsequent analyses on the basis of call rate (<0.98), suspected non-European ancestry, heterozygosity, and relatedness other than dizygotic twin status. SNPs were excluded if the minor allele frequency was smaller than 0.5%, if more than 2% of genotype data were missing, or if the Hardy Weinberg *p*-value was lower than 10^-5^. Non-autosomal markers and indels were removed. Association between SNP and the platform, batch, plate or well on which samples were genotyped was calculated; SNPs with an effect *p*-value < 10^-4^ were excluded. A total sample of 10,346 samples (including 3,320 dizygotic twin pairs and 7,026 unrelated individuals), with 7,289 individuals and 559,772 SNPs genotyped on Illumina and 3,057 individuals and 635,269 SNPs genotyped on Affymetrix remained after quality control.

Genotypes from the two platforms were separately phased using EAGLE2 (Loh et al., 2016), and imputed into the Haplotype Reference Consortium (release 1.1) using the Positional Burrows-Wheeler Transform method (Durbin, 2014) through the Sanger Imputation Service (McCarthy et al., 2016). Prior to merging, we excluded variants with info <0.75 and removed non-overlapping SNPs between platforms. After merging, we tested for minor allele frequency differences between platforms and removed SNPs with an effect p-value < 10^-4^, and Hardy Weinberg p-value > 10^-5^. Using these criteria, 7,363,646 genotyped and well-imputed SNPs were retained for the analyses.

We performed principal component analysis on a subset of 39,353 common (MAF > 5%), perfectly imputed (info = 1) autosomal SNPs, after stringent pruning to remove markers in

linkage disequilibrium (*r*^2^ > 0.1) and excluding high linkage disequilibrium genomic regions so as to ensure that only genome-wide effects were detected.

**References**

Chang, C. C., Chow, C. C., Tellier, L. C., Vattikuti, S., Purcell, S. M., & Lee, J. J. (2015). Second-generation PLINK: rising to the challenge of larger and richer datasets. GigaScience, 4(1), 7. http://doi.org/10.1186/s13742-015-0047-8

Durbin, R. (2014). Efficient haplotype matching and storage using the positional Burrows–Wheeler transform (PBWT). Bioinformatics, 30(9), 1266–1272. http://doi.org/10.1093/bioinformatics/btu014

Li, H. (2011). A statistical framework for SNP calling, mutation discovery, association mapping and population genetical parameter estimation from sequencing data. Bioinformatics, 27(21), 2987–2993. http://doi.org/10.1093/bioinformatics/btr509

Loh, P.-R., Danecek, P., Palamara, P. F., Fuchsberger, C., A Reshef, Y., K Finucane, H., et al. (2016). Reference-based phasing using the Haplotype Reference Consortium panel. Nature Genetics, 48(11), 1443–1448. http://doi.org/10.1038/ng.3679

McCarthy, S., Das, S., Kretzschmar, W., Delaneau, O., Wood, A. R., Teumer, A., et al. (2016). A reference panel of 64,976 haplotypes for genotype imputation. Nature Genetics, 48(10), 1279–1283. http://doi.org/10.1038/ng.3643

Patterson, N., Price, A. L., & Reich, D. (2006). Population structure and eigenanalysis. PLOS Genetics, 2(12), e190. http://doi.org/10.1371/journal.pgen.0020190

Price, A. L., Patterson, N. J., Plenge, R. M., Weinblatt, M. E., Shadick, N. A., & Reich, D. (2006). Principal components analysis corrects for stratification in genome-wide association studies. Nature Genetics, 38(8), 904–909. http://doi.org/10.1038/ng1847

Purcell, S., Neale, B., Todd-Brown, K., Thomas, L., Ferreira, M. A. R., Bender, D., et al. (2007). PLINK: A Tool Set for Whole-Genome Association and Population-Based Linkage Analyses. American Journal of Human Genetics, 81(3), 559–575. http://doi.org/10.1086/519795

R Core Team. (2017). R: A Language and Environment for Statistical Computing. R Foundation for Statistical Computing. Retrieved from https://www.r-project.org
